# Schema support for forming inferences in the human brain

**DOI:** 10.1101/2021.11.15.466820

**Authors:** J.P. Paulus, C. Vignali, M.N. Coutanche

**Affiliations:** Department of Psychology, University of Pittsburgh, Pittsburgh PA; Learning Research and Development Center, University of Pittsburgh, Pittsburgh PA; Center for the Neural Basis of Cognition, Pittsburgh PA

## Abstract

Associative inference, the process of drawing novel links between existing knowledge to rapidly integrate associated information, is supported by the hippocampus and neocortex. Within the neocortex, the medial prefrontal cortex (mPFC) has been implicated in the rapid cortical learning of new information that is congruent with an existing framework of knowledge, or schema. How the brain integrates associations to form inferences, specifically how inferences are represented, is not well understood. In this study, we investigate how the brain uses schemas to facilitate memory integration in an associative inference paradigm (A-B-C-D). We conducted two event-related fMRI experiments in which participants retrieved previously learned direct (AB, BC, CD) and inferred (AC, AD) associations between word pairs for items that are schema congruent or incongruent. Additionally, we investigated how two factors known to affect memory, a delay with sleep, and reward, modulate the neural integration of associations within, and between, schema. Schema congruency was found to benefit the integration of associates, but only when retrieval immediately follows learning. RSA revealed that neural patterns of inferred pairs (AC) in the PHc, mPFC, and posHPC were more similar to their constituents (AB and BC) when the items were schema congruent, suggesting that schema facilitates the assimilation of paired items into a single inferred unit containing all associated elements. Furthermore, a delay with sleep, but not reward, impacted the assimilation of inferred pairs. Our findings reveal that the neural representations of overlapping associations are integrated into novel representations through the support of memory schema.

**Significance Statement:** Our ability to draw novel links between pieces of existing knowledge allows us to understand how information in memory is related. Existing knowledge (memory ‘schema’) can facilitate learning, and then integration, of new related information. We ask how the human brain uses schema to form links between related pieces of information, and how sleep and reward affect this process. Our results suggest that memory schema helps pieces of knowledge become a single inferred unit in the brain’s memory systems. A delay with sleep between learning and retrieval, though not reward, is important for how schema achieve this.

## Introduction

The brain’s ability to process and store the information it receives allows us to accumulate knowledge that informs our immediate and future behavior. To use this knowledge flexibly, we must often draw novel links between existing pieces of knowledge. For instance, learning that Boston is North of New York, and that New York is North of Philadelphia, allows us to know that Boston is North of Philadelphia, without needing it to be explicitly taught. Such “associative inferences” (Eichenbaum, 2000; Gluck & Myers, 1993) are possibly supported by the hippocampus (Greene et al., 2006; Heckers et al., 2004) and neocortical regions (Schlichting & Preston, 2015; Spalding et al., 2018). The integration of new information into memory is supported by having a framework of related knowledge, or ‘memory schema’ (e.g., Bruett et al., 2018). The existence of a memory schema can promote rapid cortical learning (Tse et al., 2007; McClelland, 2013), likely supported by the medial prefrontal cortex (mPFC; van Kesteren et al., 2010a, 2010b, 2013; Bein et al., 2014; Brod et al., 2015; Reggev et al., 2016; van der Linden et al., 2017).

In this study, we ask how the brain uses schema to successfully infer relationships across items. During an fMRI scan, participants retrieved word pairs (half schema-congruent, half schema-incongruent) that had been directly learned, or had to be inferred. As inferred pairs are successfully integrated into memory, their neural patterns will overlap with the neural representations of the already learned underlying word pairs. If schemas draw on shared preexisting knowledge, congruent pairs should show evidence of greater memory integration, especially in cortical regions.

A second experiment examines how memory integration is modulated by two important factors: sleep (Ren & Coutanche, 2021; Durrant et al., 2015; Tse et al., 2007; Lewis & Durrant, 2011) and reward (Al-Imari & Gerlai, 2008; Wolosin et al., 2012). Based on observed neuronal replay during sleep, believed to help transfer information from hippocampal to neocortical regions (Bendor & Wilson 2012; Born et al., 2006; Walker, 2008), we hypothesized that a delay with sleep between encoding and retrieval would lead to relatively stronger memory performance and neural integration for schema-driven inferences, relative to schema-incongruent pairs.

Additionally, because dopaminergic pathways increase hippocampal activity to promote encoding (Bethus et al., 2010; O’Carroll et al., 2006), we hypothesized that reward would enhance memory consolidation, particularly through a hippocampal-mPFC circuit (Preston & Eichenbaum, 2013; Schlichting & Preston, 2016). The role of schema in reward is relatively understudied, so no prior hypotheses exist regarding the role played by reward in modifying schema-driven inferences. Either way, the findings will therefore establish an understanding of their relationship.

## Methods

### Participants

Participants were native English speakers with normal or corrected-to-normal vision and no learning or attention disorders.

For experiment 1, participants were recruited until 22 usable participants’ datasets were available for analysis. Data from 28 participants were collected to achieve the 22-subject number, after excluding 6 from analyses based on their poor behavioral accuracy (< 60% on the immediate recognition task for directly learned pairs). The remaining 22 participants (16 females; mean [*M*] age = 23; standard deviation (SD) = 2.9) were included in all analyses.

In experiment 2, data were collected from an independent sample of 20 healthy adults (15 females; *M* age = 22.3; SD = 3.1). Following the emergence of COVID-19 cases and the resulting changes in regulations imposed by the University of Pittsburgh, data collection was interrupted before reaching the goal of 22 participants. Four individuals were excluded from analyses due to poor retrieval accuracy (<60%) and one individual was excluded due to missing anatomical data, resulting in 15 suitable participants for further analysis.

Participants were compensated for their participation in the study. All procedures were approved by the Institutional Review Board of the University of Pittsburgh.

### Study Materials

To study the effect of schema congruency on the retrieval of word pairs that were directly or indirectly associated in the learning phase, we used a 2 (schema congruent vs. schema incongruent) x 2 (directly learned vs. inferred) within-subjects study design. We adapted and expanded 28 sets of words (32 sets for experiment 2) from Zhang et al. (2018), with each set containing four nouns (E.g., surgeon - school - sailor - cottage). In line with previous memory studies investigating associative inference (van der Linden et al., 2017; Shohamy & Wagner, 2008; Spalding et al., 2018; Zhang et al., 2018), each of the four concepts in a set represents a component of the AB-AC inference paradigm (A - B - C - D). This design allows us to test a participant’s memory of the word pairs that were directly learned in the learning phase (AB, BC, CD; direct pairs), as well as word pairs that were indirectly learned (associated by several links: AC, AD; inferred pairs). In half of the sets, the four words were strongly thematically related within a schema (e.g., teacher - classroom - student - dormitory), while the remaining sets were incongruent to a single schema (e.g., surgeon - school - sailor - cottage). A and C represented people, and B and D represented locations. For experiment 2, two additional sets were added to the congruent and incongruent conditions, resulting in 32 total sets (16 congruent, 16 incongruent), in order to increase statistical power for the additional reward factor (half associated with a reward; details below).

Words within a congruent set were selected based on their semantic relatedness; calculated through Latent Semantic Analysis (LSA; Landauer & Dumais, 1997). LSA can be used to measure the semantic association between two words by observing their concurrence in a large corpus of texts (Landauer & Dumais, 1997). Higher LSA values correlate with a stronger semantic relationship between the two words. Given the nature of our forced-choice training task, each set additionally had six location foils and three people foils that were either congruent or incongruent to the schema, depending on the congruence condition of the set. While the foils for the incongruent sets did not share a schema with that set’s A, B, C, or D, they shared a schema with words seen in one of the congruent sets. This design prevented participants from selecting the correct pairs of words during the forced-choice tasks simply based on their knowledge that the foil did not belong to any schema they had encountered during the training.

### Experimental Design

#### Overview

The experimental procedure was amended from a recent behavioral study of memory performance based on schema congruency (Zhang et al., 2018). All task procedures were created in Matlab 2017a using the Psychophysics Toolbox Version 3 (PTB-3) package for presenting stimuli on-screen. Prior to any participation, participants were taken through the informed consent process, after which they completed a brief demographic questionnaire. Participants were initially trained on a computer to learn word pairs (AB, BC, CD) using a forced-choice associative inference task until they successfully reached criterion (more details below; Fig. 1A). Memory for the word pairs that were explicitly taught (AB, BC, CD) and (untaught) inferred pairs (A-C, A-D) were subsequently tested in a similar task (Fig. 1C). Following the retrieval task, participants performed a localizer in which they rated how likely two objects (belonging to one of the schemas from the study) are found in the same location (experiment 1) or a monetary incentive delay (MID) task (adapted from Knutson et al., 2001). These localizers were not used in analyses of this experiment. After the scan, participants completed several memory and sleep related self-report questionnaires.

**Figure 1.**
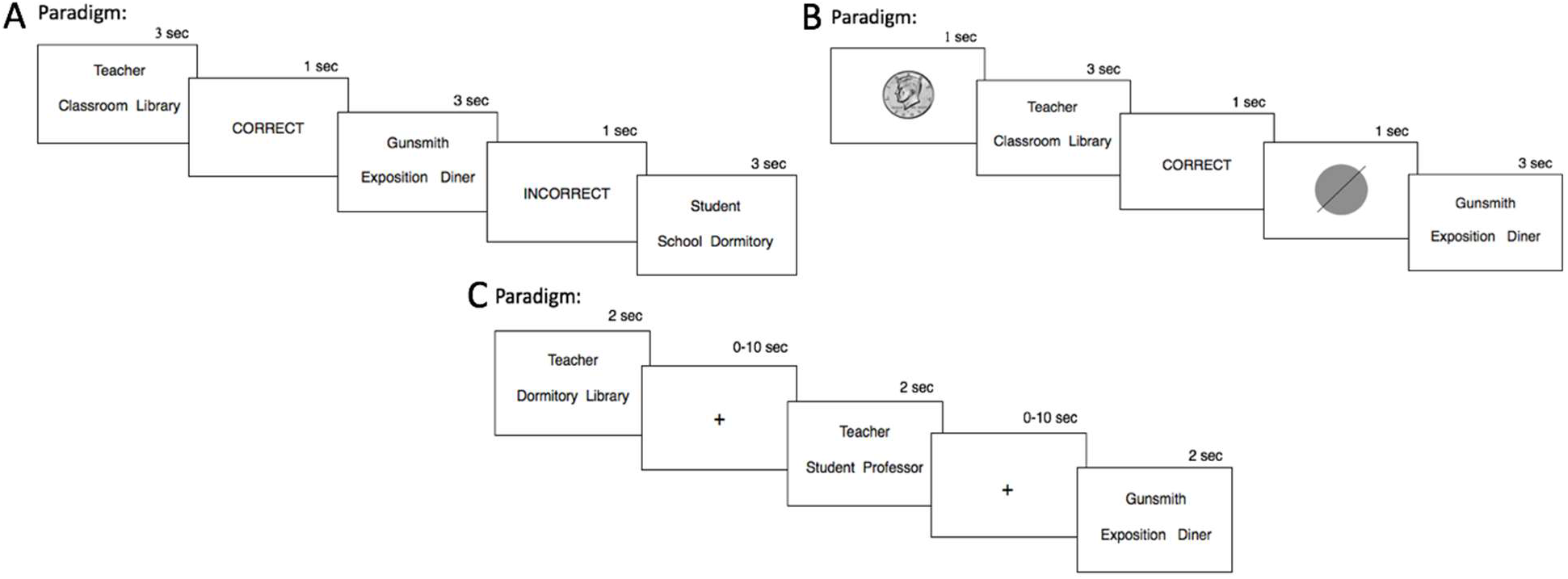
Experimental procedure for the training phase in experiment one (A) and experiment two (B), as well as the retrieval phase for both experiments (C).

**Figure 2.**
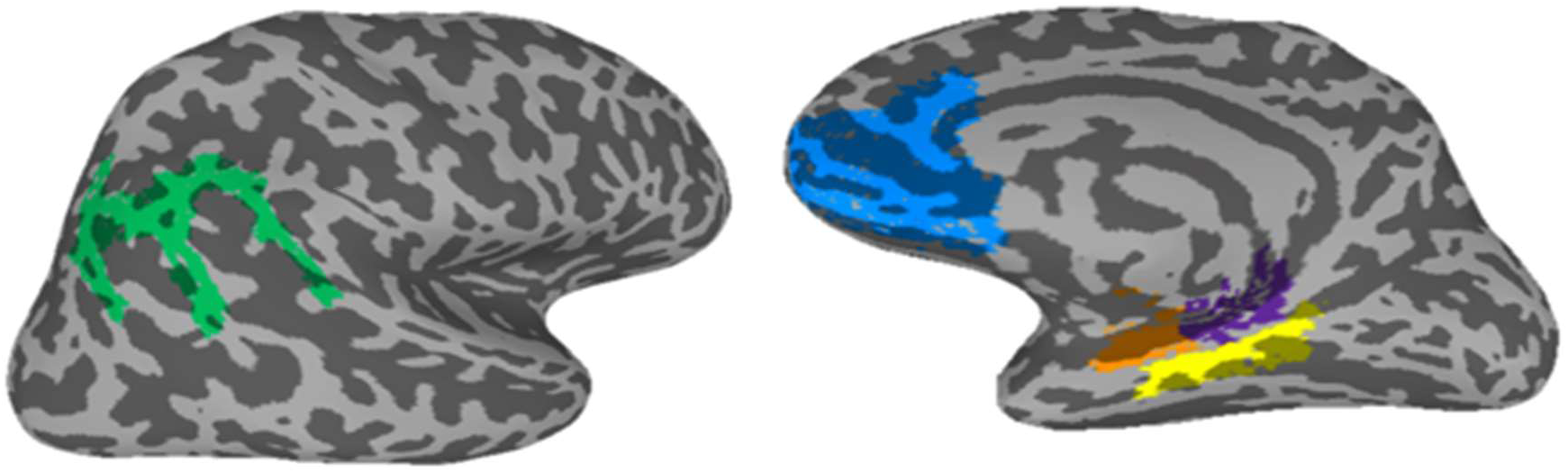
The anatomically-defined regions of interest (PHc: Yellow; AG: Green; mPFC: Blue; antPHC: Orange; posHPC: Purple) used in the current study.

#### Training Phase

During experiment 1, participants were trained to learn associated word pairs through a forced-choice paradigm, though they were never explicitly told that they would be subsequently tested on their memory of the word pairs. Participants were presented with a cue word (representing A, B, or C from a set) at the top of the screen. Two additional words, the associated word (target) and a foil, were shown on the bottom left and right of the screen (3 sec). The positions for target items and foils for each presentation were randomized so that the target item was presented equally on each side of the screen across trials. Participants were instructed that they had three seconds to select the word that was associated with the cue. Following the participant’s response, “Correct” or “Incorrect” was displayed in the center of the screen for 1 second to indicate whether they had selected the correct word. If no response was recorded in the allotted 3 seconds, the screen displayed “Too Late” and proceeded to the next item, without showing the correct choice. As participants were blind to the correct associations on their first encounter with a certain cue word, they would have to randomly choose one of the two choice words (50% chance of successfully selecting the target word) and rely on feedback (1 sec) to learn and inform their response for subsequent trials.

The task was divided into four sections, each containing a different subset of the word pairs. Once all of the words within a section were shown once, the word pairs were shown again in the same order to maximize the amount of time between presentations. In all repeat instances of a given item, the foil was always the same. Pairs were removed from the pool after three consecutive correct trials, thereby reducing the number of pairs presented until all pairs were learned to this criterion. Once all four sections were completed, the entire process was repeated for two more cycles to ensure that the word pairs had been successfully learned. While the cue word was paired with the same target word across the three cycles, the foils were unique to that cycle.

Experiment 2 followed the same training protocol with the addition of reward cues. To investigate the influence of reward, certain word pairs were associated with a possible future monetary reward. Participants were instructed that correctly learning rewarded items would lead to greater reward later in the study. A symbol indicating the reward potential for each item was presented at the center of the screen before every word pair (1 sec). A picture of a half dollar coin was used to indicate that the following item would be rewarded, while unrewarded items were preceded by a gray circle of the same size with a line through it (Fig. 1B). Due to the increase in sets from experiment 1, the training was divided into 8 sections (rather than the 4 in Experiment 1).

#### Retrieval Phase

The retrieval phase took place immediately after the training phase (experiment 1) or approximately 24 hours later (experiment 2). Although the retrieval phase was designed to be as similar to the training phase as possible, some changes were made to adapt the task for fMRI. A rapid event-related (rER) design was used, with OptSeq2 providing the presentation sequence to optimize extraction of the signal (Dale, 1999). Within stimulus presentations, a jittered fixation cross (0-10 sec) was shown to provide a buffer between groups of stimuli, allowing us to isolate the hemodynamic response function associated with each stimuli type.

In the retrieval task, participants were tested on their memory of the word pairs they had learned in the training. As in the training, the cue word was displayed at the top of the screen.

Participants selected which of the two words on the bottom of the screen had been associated with the cue during training, using two buttons on an fMRI compatible response glove. The location of the target word and foil was randomized to appear on the left and right side of the screen equally. Unlike the training task, participants only had 2 seconds (from 3 secs in training) to respond before the trial was marked as incorrect. Participants were instructed to respond as quickly and as accurately as possible and that no feedback would be provided. In addition to being tested on the directly learned pairs from the training (AB, BC, CD), participants would sometimes have to select target words that were indirectly associated (inferred; AC, AD). The order of presentation was randomized across runs; however, inferred pairs were always presented before direct pairs so that retrieval of direct pairs would not interfere by reinforcing these relations. The retrieval task was separated into four functional runs. The first run contained half of the word pairs, and the second run contained the other half. Runs 3 and 4 consisted of the word pairs from runs 1 and 2 respectively, but with different foils (selected from the training phase), so that each word pair was tested twice in the retrieval phase. Each run had an equal number of schema congruent and incongruent trials.

In experiment 2, the items that had been associated with a reward cue during the training were now eligible for additional monetary compensation based on accuracy on those items.

Participants were instructed that each rewarded pair would be associated with a $0.50 reward. Since we were interested in the effect of reward during learning, the reward status of each item (previously reward-cued versus not) was not indicated during the recognition task.

### ROI Creation

We used Freesurfer to create anatomically-defined regions of interest (ROIs) for each participant. We created ROIs for the mPFC, a neocortical region found to be involved in the processing of schema congruent information, as well as the hippocampus (van Kesteren et al., 2010, 2012; Spalding et al., 2018; Schlichting & Preston, 2014). An additional neocortical region, the angular gyrus (AG), implicated in schematic memory was included in our ROI selection (Gilboa & Marlatte, 2017; Wagner et al., 2015). Following literature that has found functional differences between the anterior and posterior hippocampus (Bein et al., 2019), we manually divided the hippocampus into two halves along the coronal axis using AFNI (Cox, 1996). To investigate the role of reward on rapid memory integration, we created ROIs for the nucleus accumbens (NAc) and substantia nigra (SN; Knutson et al., 2001; Adcock et al., 2006).

### fMRI Data

#### fMRI acquisition

The scanning parameters were identical for both experiments. Data were collected on a 3.0T Siemens Magnetom Verio equipped with a 32-channel coil. For blood-oxygen level dependent (BOLD) fMRI images, T2-weighted scans were acquired using the following parameters: repetition time (TR) = 2000 ms, echo time (TE) = 30 ms, 2mm slice thickness with 69 slices per run, 2×2×2 mm voxel size, field of view = 212×212 mm, flip angle = 79°. For anatomical images, taken after functional scans, T1-weighted scans were taken from the sagittal plane at 1mm isotropic resolution using an MPRAGE scan with: TR = 2300 ms, TE = 1.97 ms, inversion time (TI) = 900 ms, slice thickness = 1 mm, resolution size = 1 mm isotropic, flip angle = 9 degrees, field of view = 256×256 mm.

#### Preprocessing

All preprocessing was conducted using AFNI (Cox, 1996). Functional data were slice-time corrected, aligned to a base volume, and scaled to have a mean signal of 100 with lower and upper bounds of 0 and 200. Visual inspection of the motion time series confirmed that all participants moved no more than 1.5 mm once in any dimension from one volume to the next.

#### MVPA Decoding

Multi-voxel pattern analysis (MVPA) was conducted using a two-fold cross-validation structure, in order to investigate the information within patterns of activation for directly learned and inferred associations within ROIs. For a given condition, beta coefficients for each item in runs 1 and 3, and then runs 2 and 4, were determined using a generalized linear model. The beta coefficients were then used to train a Gaussian Naïve Bayes (GNB) model. To balance the number of trials from each condition, a randomized subset of directly learned trials were selected to match the smaller number of inferred trials. This process was repeated 100 times in both directions (training on items from runs 1 and 3, and then on items from runs 2 and 4) for each ROI and participant. Accuracy for the model was calculated for each fold within an iteration, and averaged after completion of all 100 iterations. Group-level tests determined if classifier accuracy across participants differed significantly from chance.

## Results

### Experiment 1

#### Behavioral Results

Six participants were removed due to low retrieval accuracy (<60%) on the directly learned pairs. The remaining 22 participants demonstrated above-chance retrieval performance for direct (p < .001, t(21) = 17.92) and inferred (p = .008, t(21) = 2.92) pairs.

As expected, participants performed significantly better (p < .001, t(21) = 5.86) on word pairs they had directly learned (M = .83; sd = .09) than on inferred pairs (M = .61; sd = .18), including for schema-congruent (p < .001, t(21) = 5.59) and incongruent (p < .001, t(21) = 5.68) separately (Fig. 3). The direct and inferred pairs showed a significant difference in response-times (p < .001, t(21) = -7.70) with faster RTs for direct (M = 1.29; sd = .11) than inferred (M = 1.49; sd = .17), in both the congruent (p < .001, t(21) = -7.03) and incongruent (p < .001, t(21) = -7.33) conditions. For more details on behavioral performance, see Table 2.

**Figure 3.**
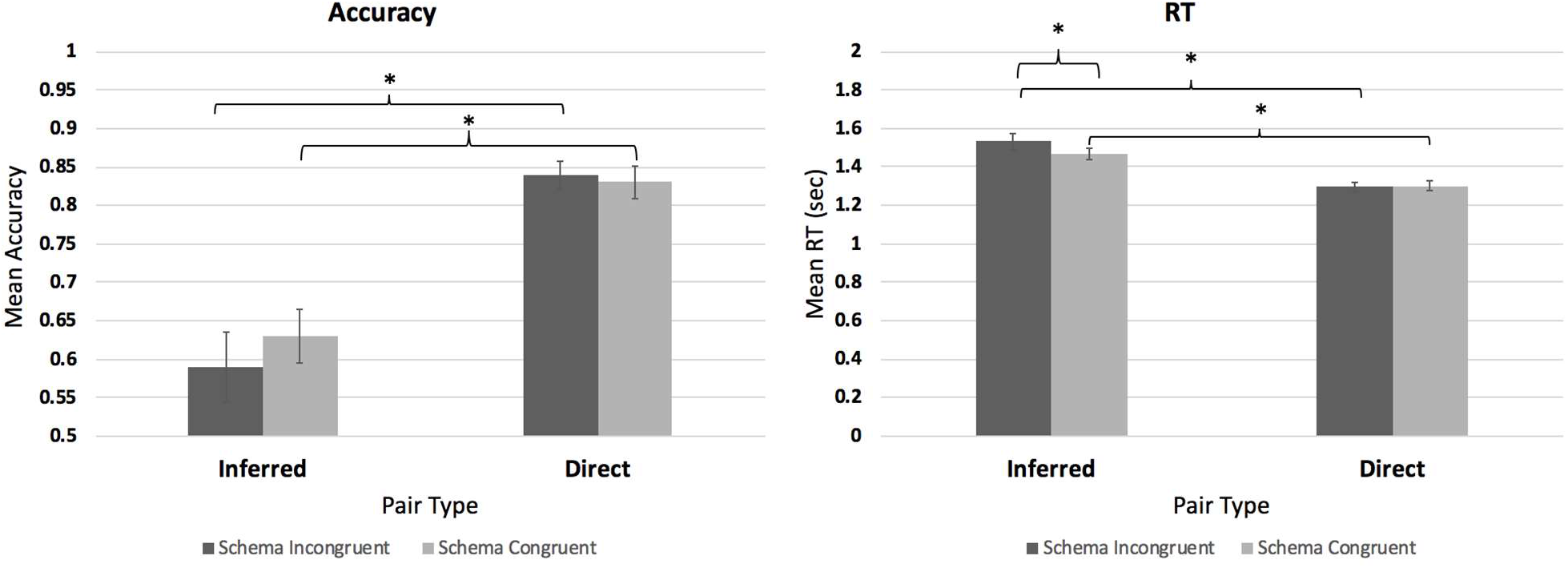
Mean Accuracy and Response Time (RT) between inferred and direct pairs based on schema consistency. Error bars reflect standard errors of the mean.

**Table 1.** Example Sets of Schema Congruent and Schema Incongruent Words.

| Condition | Stimulus Type |  |  |  |
| --- | --- | --- | --- | --- |
|  | *Person 1* | *Location 1* | *Person 2* | *Location 2* |
| Schema Congruent | Teacher | Classroom | Student | Dormitory |
| Schema Incongruent | Baker | Mountain | Principal | Circus |

**Table 2.** Behavioral Performance on Retrieval Task of Experiment 1.

|  | Direct | Inferred | Collapsed |
| --- | --- | --- | --- |
| Congruent | 0.83 (0.10) | 0.63 (0.16) | 0.75 (0.10) |
| Incongruent | 0.84 (0.08) | 0.59 (0.21) | 0.74 (0.11) |
| Collapsed | 0.83 (0.09) | 0.61 (0.18) | 0.74 (0.10) |

In order to check whether rapid memory integration was successful in the schema congruent and incongruent items, we tested whether group accuracies in the inferred pairs exceeded chance. Both the congruent inferred (p = .001, t(21) = 3.74) and incongruent inferred (p = .047, t(21) = 2.11) pairs were significantly above chance. Accuracy did not differ between congruent and incongruent inferred pairs (p = .171, t(21) = 1.42). Incongruent inferred RTs were significantly slower than congruent inferred RTs (p < .001, t(21) = -3.89).

#### fMRI Results

##### Distinctions between directly learned and inferred associations

We first trained a GNB classifier to distinguish between directly learned versus inferred pairs. We found significant classification (Fig. 4) of these types of pairs in the AG (p < .001, t(21) = 4.29), mPFC (p = .004, t(21) = 3.25), and posterior hippocampus (p = .046, t(21) = 2.12), but not in the PHc (p = .240, t(21) = 1.21) or anterior hippocampus (p = .880, t(21) = .153). After restricting the classification to only the schema congruent items (Fig. 5), the AG (p = .003, t(21) = 3.34) and mPFC (p < .001, t(21) = 3.93) showed above-chance classification performance, with a trending result in posterior hippocampus (p = .062, t(21) = 1.97), and no significance in the PHc (p = .205, t(21) = 1.31) or anterior hippocampus (p = .957, t(21) = .055). An analogous classification in the incongruent pairs revealed above-chance classification in the AG (p = .002, t(19) = 3.60), with trending significance in the mPFC (p = .091, t(19) = 1.78) and posterior hippocampus (p = .079, t(19) = 1.86). No significant classification performance was found in the PHc (p = .980, t(19) = -.026) or anterior hippocampus (p = .957, t(19) = -.201).

**Figure 4.**
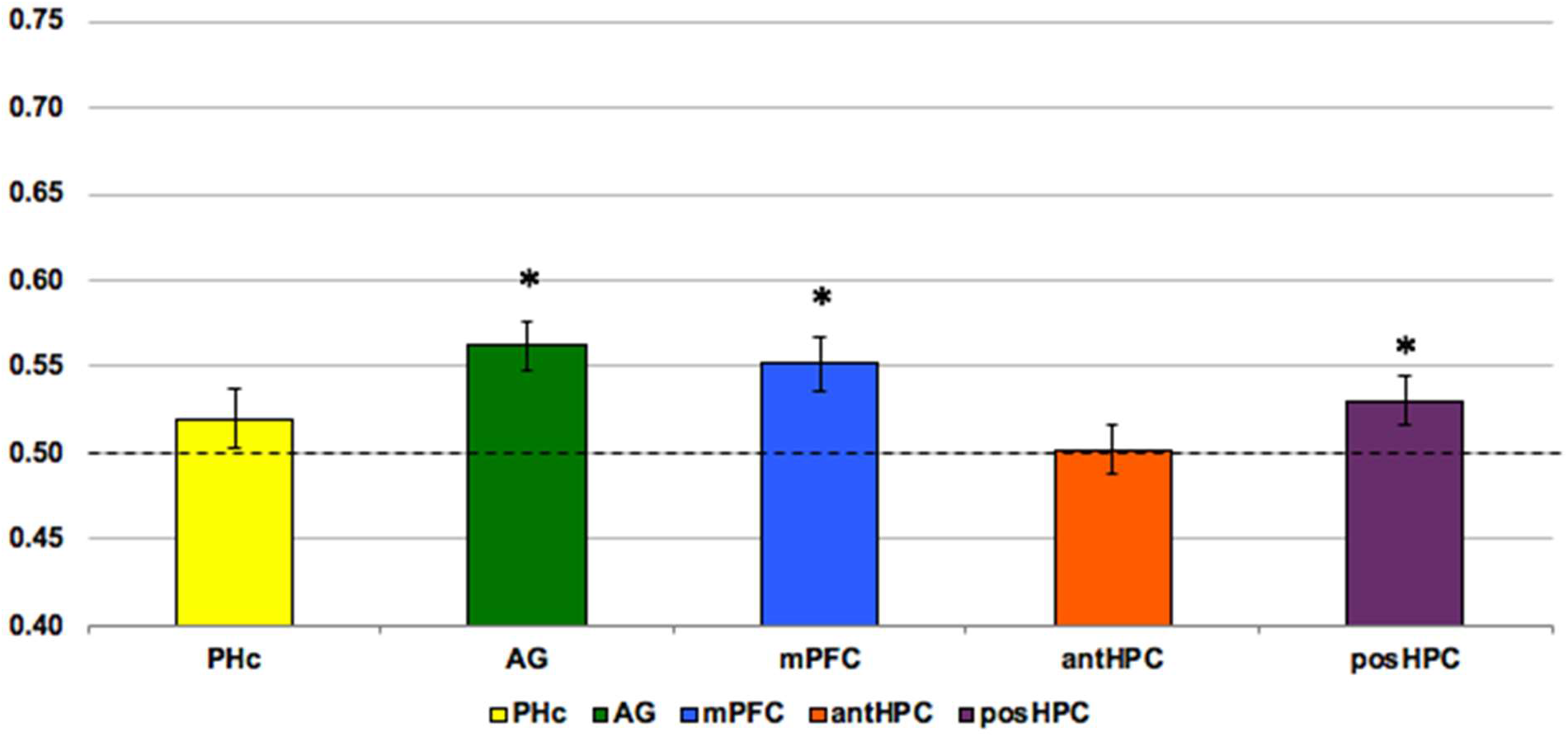
MVPA classifier accuracies for retrieval of directly learned vs. inferred pairs within all sets (^*^ indicates significance; --- indicates chance). Error bars reflect standard error of the mean. Colors correspond to Figure 2.

**Figure 5.**
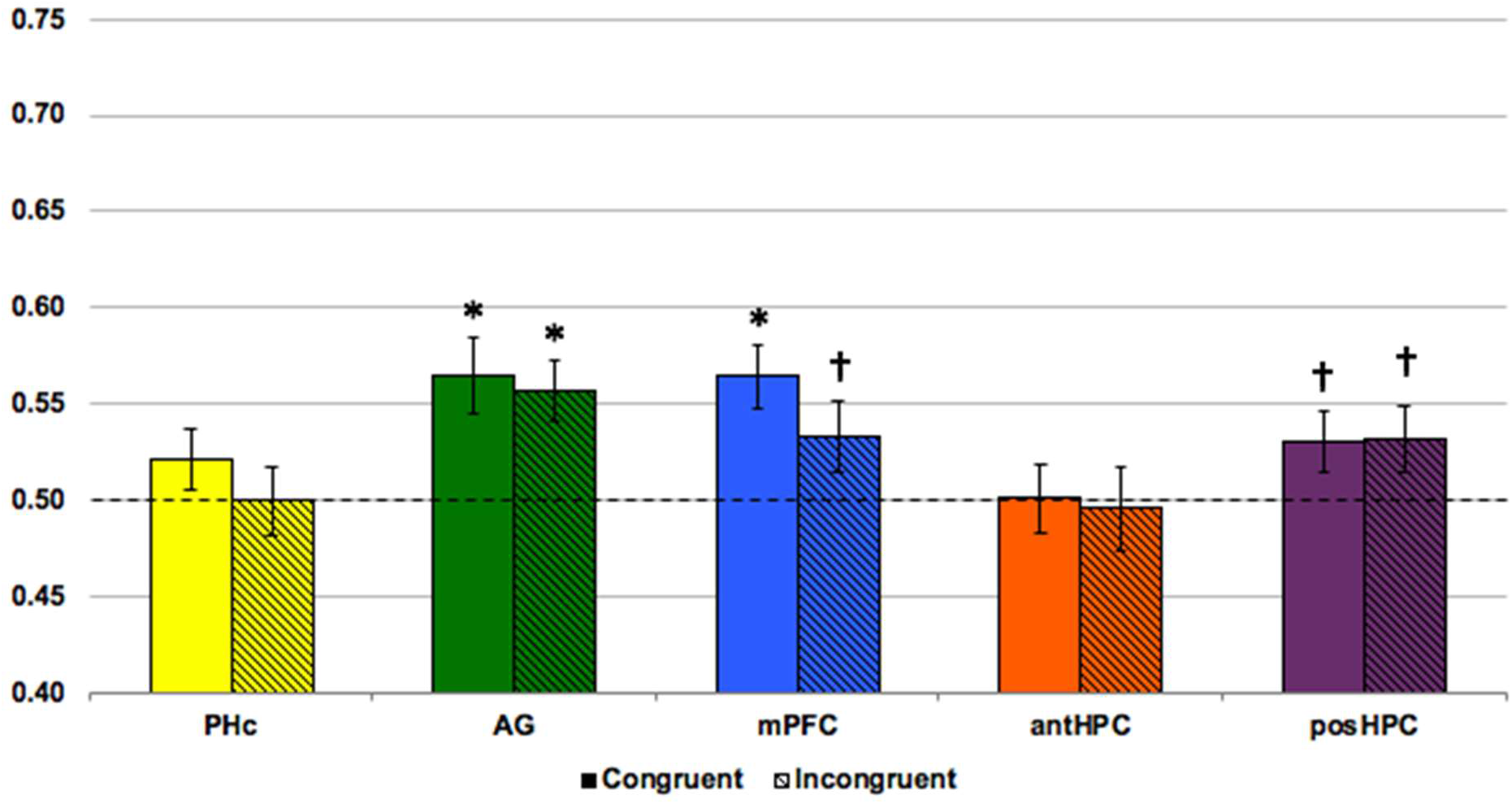
MVPA classifier accuracies for retrieval of directly learned vs. inferred pairs within congruent and incongruent items (^*^ indicates significance; † indicates trending significance;---indicates chance). Error bars reflect standard error of the mean. Colors correspond to Figure 2.

##### Reactivation of direct associations while retrieving inferred pairings

We next examined whether neural patterns for the directly learned AB and BC pairs were reactivated during the retrieval of the linked AC pairs. We correlated the patterns for each set’s AB pair with the same set’s AC pair. Similarly, we also correlated the BC pair with the AC pair. In a linear mixed effects (LME) regression model for each ROI, schema-congruency was a significant predictor (Fig. 6A) of the correlation between the inferred pair (AC) and its constituents (AB and BC) in the PHc (B= -.023, p=.025), mPFC (B=-.028, p=.024), and posterior hippocampus (B=-.017, p=.024), with higher *r*-values for congruent pairs (M_PHc_ = .0347; M_mPFC_= .0795; M_posHPC_ = .0336) than for incongruent pairs (M_PHc_ = .0115; M_mPFC_ = .0517; M_posHPC_ =.0166), but not in the anterior hippocampus (B=.010, p=.375) or AG (B=-.016, p=.221). The order of the direct pairings (AB or BC being correlated with AC) was also a significant predictor in anterior hippocampus (B=.030, p=.009), posterior hippocampus (B=-.035, p<.001), PHc (B=.037, p<.001), AG (B=.026, p=.044), and mPFC (B=.025, p=.039), suggesting that the order of the words was an important factor.

**Figure 6.**
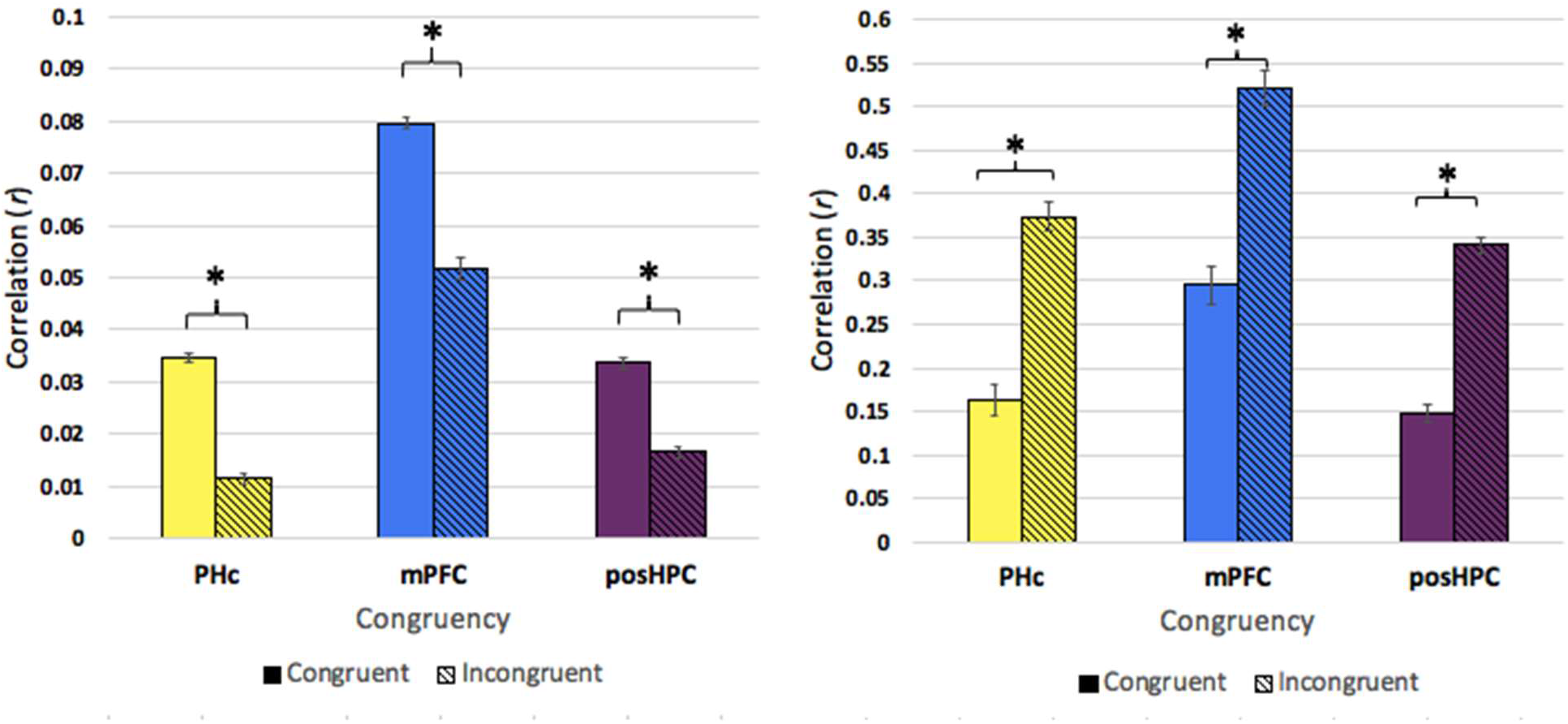
(A) Correlations of inferred pair AC with its constituents AB and BC in schema congruent and incongruent items. AC patterns were correlated with AB and BC individually, after which the resulting r values were averaged. (B) Correlations of direct pairs AB and BC by congruency, suggesting that the higher correlations of AC and its constituents are not due to shared semantic content in congruent pairs. Error bars reflect standard error of the mean. Colors correspond to Figure 2.

**Figure 7:**
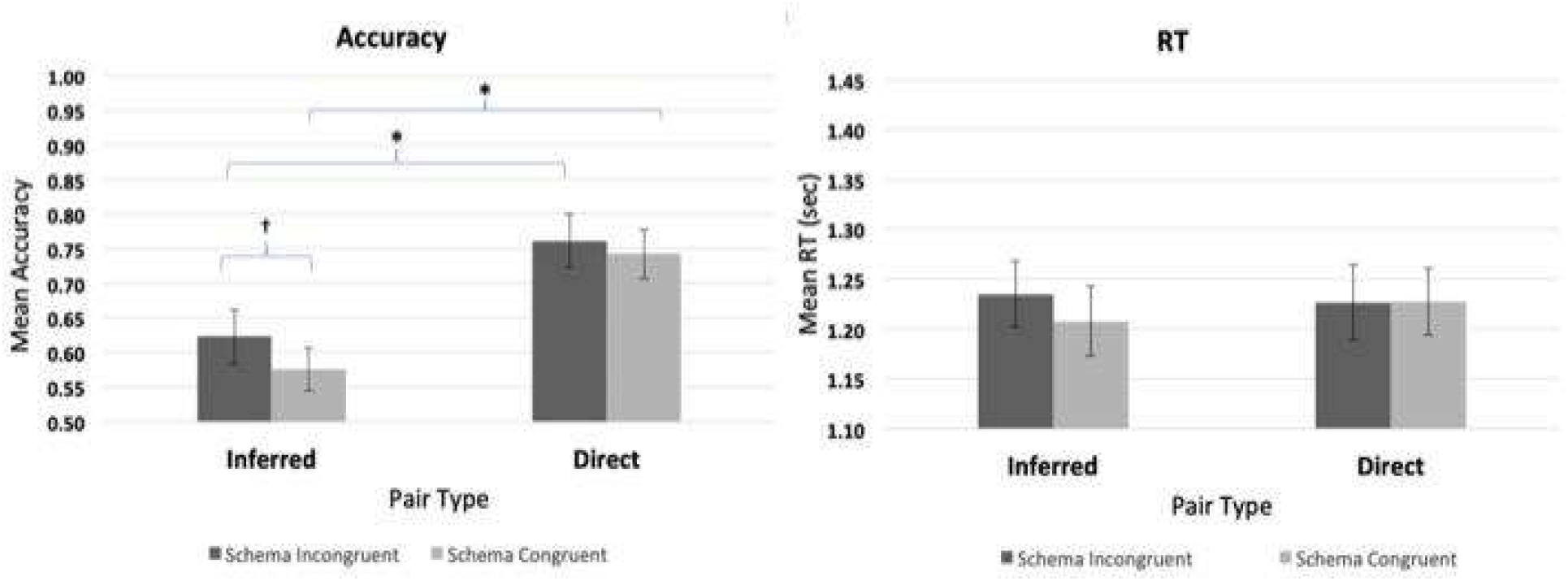
Accuracies and response times for retrieval of incongruent vs. congruent unrewarded pairs (^*^ indicates significant difference; † indicates trending significance). Error bars reflect standard error of the mean.

In order to see whether the higher correlations in congruent pairs are simply a result of semantic similarity within the same schema (e.g., due to a shared topic), we correlated AB with BC and found that incongruent sets were more highly correlated than congruent sets in all three regions (PHc: p < .001, t(21) = -10.70; mPFC: p < .001, t(21) = -10.52; posterior hippocampus: p < .001, t(21) = -16.82), suggesting that the higher pattern reactivation of the constituent pairs during retrieval is not simply due to overlapping semantic content (Fig. 6).

### Experiment 2

#### Behavioral Results

Overall, participants recorded a mean retrieval accuracy of 67.83% on all directly learned trials, indicating that items were directly learned to criterion on day one. Three-factor repeated measures analysis of variance (ANOVA) between congruency, reward, and inference indicated no main effect of congruency on retrieval performance (F(1,14) = 0.012, p = 0.92). However, the same analysis showed a significant main effect of inference on retrieval accuracy (F(1,14) = 185.264, p < .001) as well as a significant main effect of reward (F(1,14) = 6.017, p = 0.028). No significant interaction was observed between any of our three factors.

No differences were observed in retrieval performance between congruent and incongruent directly learned items during retrieval (p = 0.991, t(14) = 0.012), which suggests that any difference between congruent and incongruent accuracies found elsewhere are not due to differing degrees of learning. Significant differences were found between rewarded and unrewarded retrieval accuracies (p = 0.010, t(14) = -3.005, Table 3), however the unrewarded group (M = 0.736) outperformed the rewarded group (M = 0.699), which was not expected.

**Table 3:** Mean behavioral accuracies by condition. Standard deviations are shown in parentheses. Note that due to different numbers of stimuli in conditions (e.g. more direct than inferred pairs), a given mean might be more affected by one condition than another.

| **Within Unrewarded** | Direct | Inferred | Collapsed |
| --- | --- | --- | --- |
| Congruent | 0.80 (0.11) | 0.64 (0.12) | 0.74 (0.10) |
| Incongruent | 0.78 (0.13) | 0.60 (0.13) | 0.72 (0.12) |
| Collapsed | 0.79 (0.11) | 0.63 (0.09) | 0.73 (0.09) |
| **Within Rewarded** | Direct | Inferred | Collapsed |
| Congruent | 0.74 (0.15) | 0.58 (0.12) | 0.68 (0.13) |
| Incongruent | 0.76 (0.14) | 0.62 (0.15) | 0.71 (0.13) |
| Collapsed | 0.75 (0.13) | 0.60 (0.13) | 0.69 (0.12) |

**Table 4:** Mean table of classifier accuracies averaged across ROI for experiment 1 (no sleep) and experiment 2 (sleep)

|  | No sleep | Sleep |
| --- | --- | --- |
| Congruent | 0.532 (.045) | 0.528 (.062) |
| Incongruent | 0.524 (.063) | 0.581 (.062) |

**Table 5:** Mean table for RSA correlations averaged across ROI for experiment 1 (no sleep) and experiment 2 (sleep)

|  | No sleep | Sleep |
| --- | --- | --- |
| Congruent | 0.073 (.038) | 0.131 (.035) |
| Incongruent | 0.058 (.033) | 0.067 (.035) |

Investigating this further, a main effect of reward was observed for congruent sets (F(1,14) = 5.678, p = 0.032), however no main effect of reward was seen for incongruent sets (F(1,14) = 1.792, p = 0.202). These results suggest that the effect of reward discovered initially results from its influence on congruent sets. Additionally, significant differences in accuracy were observed for rewarded congruent pairs (p = 0.018, t(14) = -2.693) but not for rewarded incongruent pairs (p = 0.123, t(14) = -1.639). When collapsing across reward, behavioral results of experiment 2 mirrored those observed in experiment 1, so motivation was collapsed for the remaining analysis to observe the effects of a delay with sleep.

Finally, congruent directly learned and inferred pairs showed a significant difference in retrieval accuracy (p < 0.001, t(14) = 9.222); this results was replicated for incongruent pairs (p < 0.001, t(14) = 8.878). Both congruent (p < 0.001, t(14) = 4.908) and incongruent (p = 0.0015, t(14) = 3.932) linked pairs were retrieved at levels above chance, which is predictive of memory integration.

Paired t-tests of subject reaction times for items showed significant differences in response time for a number of different stimuli types. As expected, response times for the retrieval of directly learned and inferred pairs were significantly different (M_RT-Direct_ = 1.228 secs, M_RT-Inferred_ = 1.328 secs, p < .001, t(14) = -6.97). Directly learned congruent items showed significantly faster response times compared to inferred congruent (p < .001, t(14) = -7.85) but was not significantly faster than directly learned incongruent items (p = .383, t(14) = -.900).

Similarly, response times for directly learned incongruent items were significantly faster than incongruent inferred (p < .001, t(14) = -5.51). No significant difference was recorded between congruent inferred and incongruent inferred items (p = .397, t(14) = -.874). In further analysis, response times were broken down into direct/inferred and then rewarded/unrewarded groups (Fig. 8). Here we found evidence that in all conditions, reward led to faster response times (p < 0.001). However when collapsing across congruency and connection, no significant difference was observed between rewarded and unrewarded items (p = 0.879, t(14) = -0.155).

**Figure 8:**
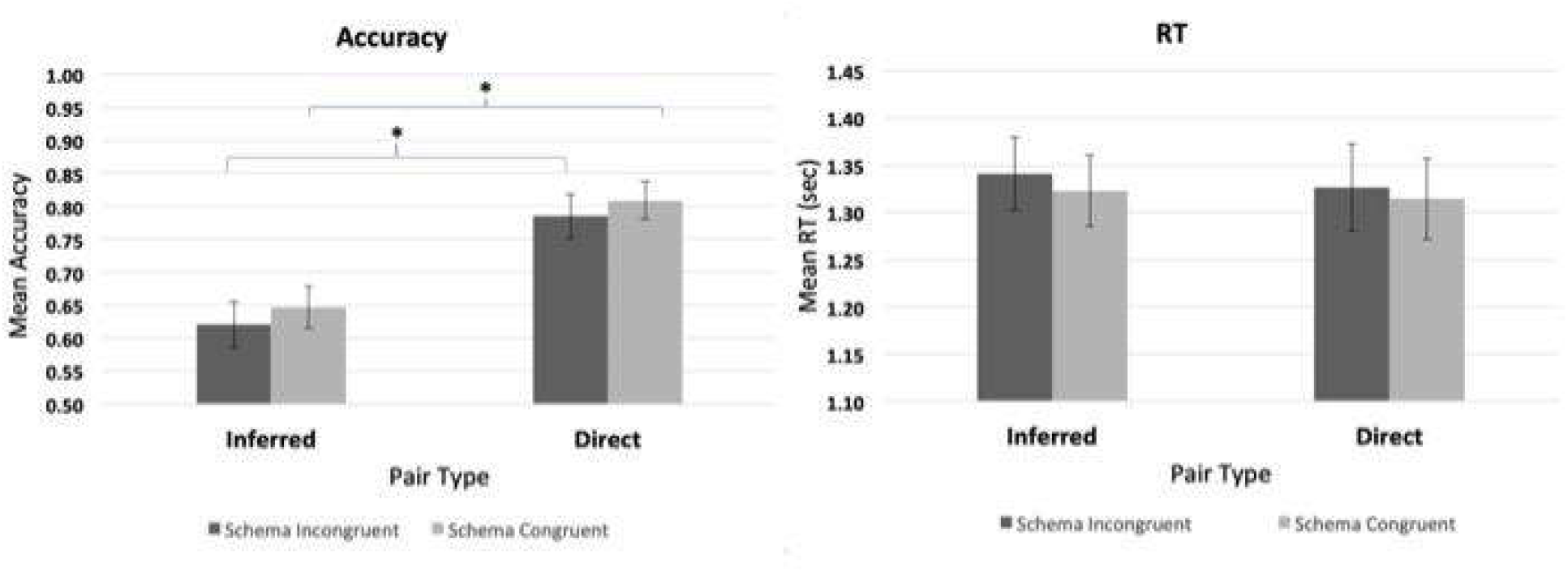
Accuracies and response times for retrieval of incongruent vs. congruent rewarded pairs (^*^ indicates significant differences). Error bars reflect standard error of the mean.

#### fMRI Results

##### MVPA Decoding

The primary analysis tested for differences in pattern activation for ROIs during retrieval of directly learned and inferred items. Here, we observed differences in patterns for inferred and directly learned items within the AG (M = .630, t(14) = 7.27, p < .001), and the mPFC (M= .572, t(14) = 4.54, p < .05), with results trending towards significance in the PHc (M = .537, t(14) = 1.81, p = .092), while the anterior (M = .512, t(14) = .821, p = .371) and posterior (M = .543, t(14) = 1.74, p = .104) hippocampal subdivisions, failed to reach significance (Fig. 10).

**Figure 9:**
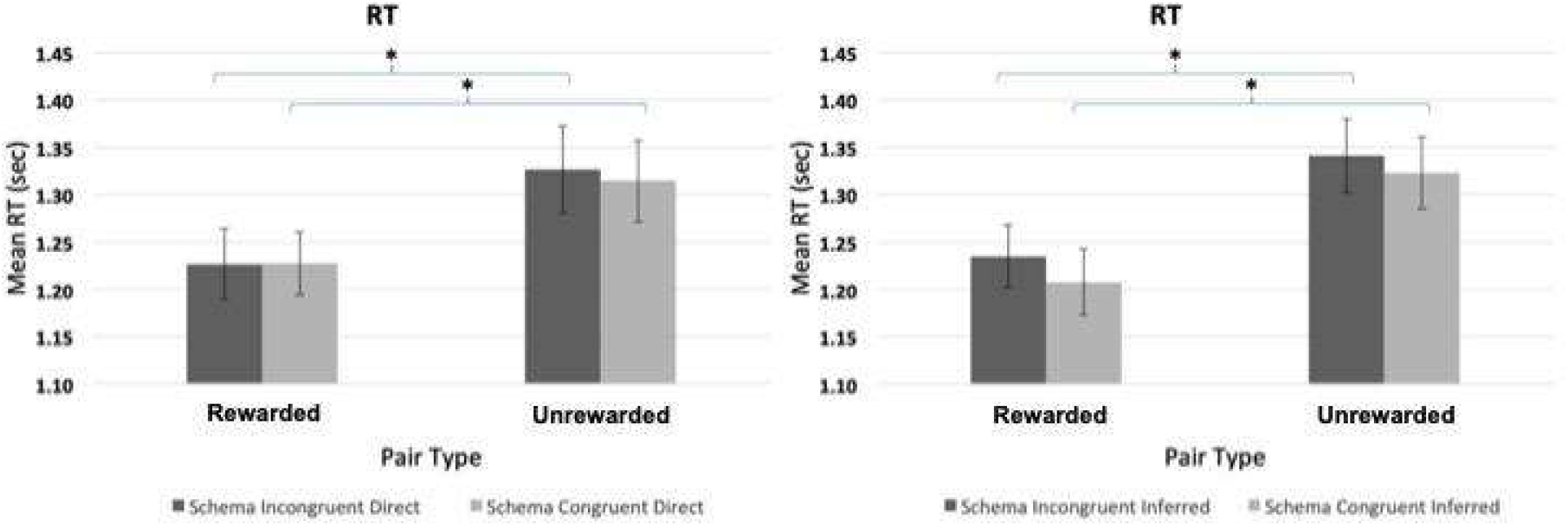
Response times for retrieval of rewarded vs. unrewarded inferred incongruent/congruent pairs (left) and direct incongruent/congruent pairs (right; ^*^ indicates significant differences).

**Figure 10:**
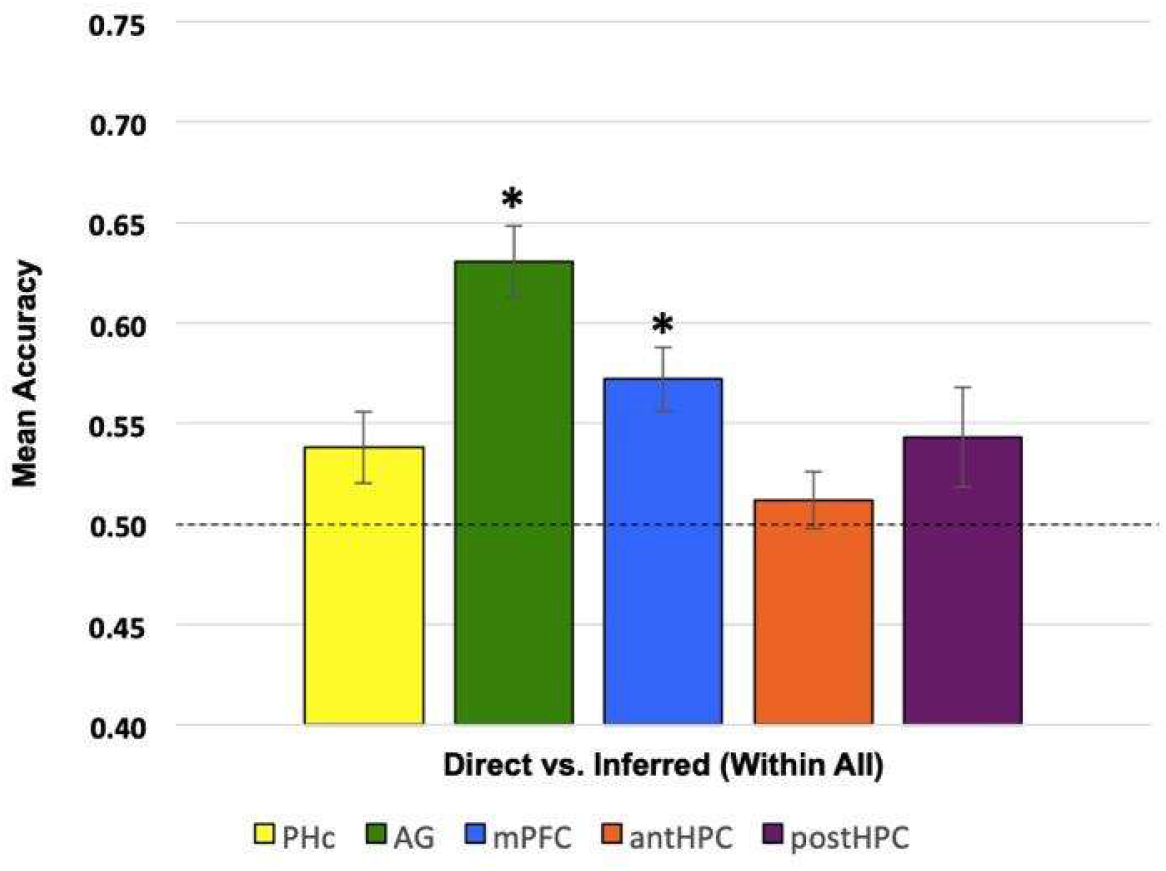
MVPA classifier accuracies for retrieval of directly learned vs. inferred pairs (^*^ indicates significance, --- indicates chance). Error bars reflect standard error of the mean. Colors correspond to Figure 2.

Next, dividing our data to analyze patterns of activation during retrieval of congruent items (congruent-inferred vs. congruent-direct) revealed significant above-chance classification within the AG (M = .558, t(14) = 2.53, p < .05), with trending results in the PHc (M = .540, t(14) = 1.87, p = .083), while no significance was found in the mPFC (M = .524, t(14) = 1.60, p = .133), anterior hippocampus (M = .503, t(14) = .121, p = .905), or posterior hippocampus (M = .519, t(14) = .823, p = .424; Fig. 11).

**Figure 11:**
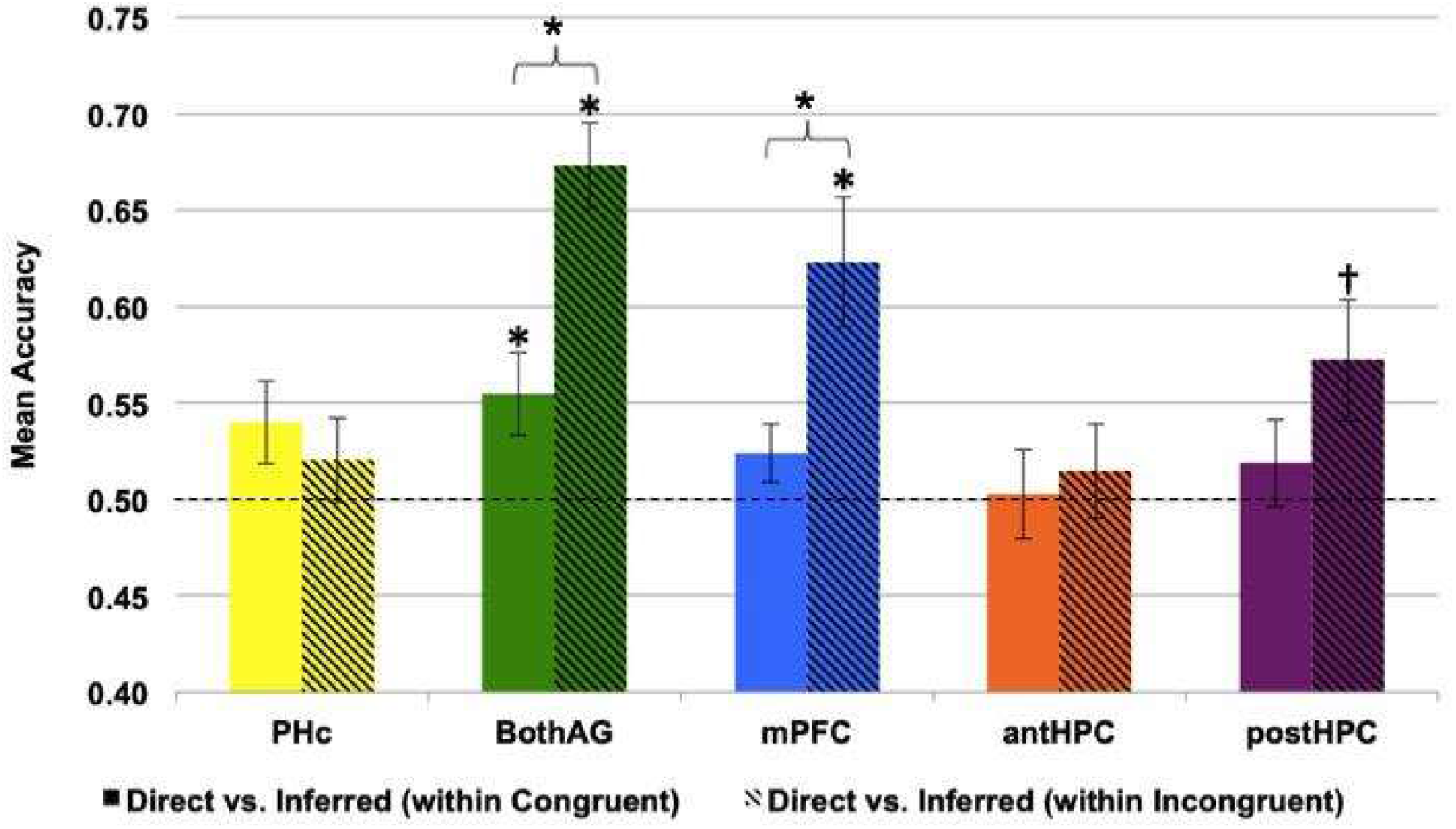
MVPA classifier accuracies for retrieval of congruent (solid) and incongruent (slashed) directly learned vs. inferred pairs (^*^ indicates significance, --- indicates chance). Error bars reflect standard error of the mean. Colors correspond to Figure 2

**Figure 12:**
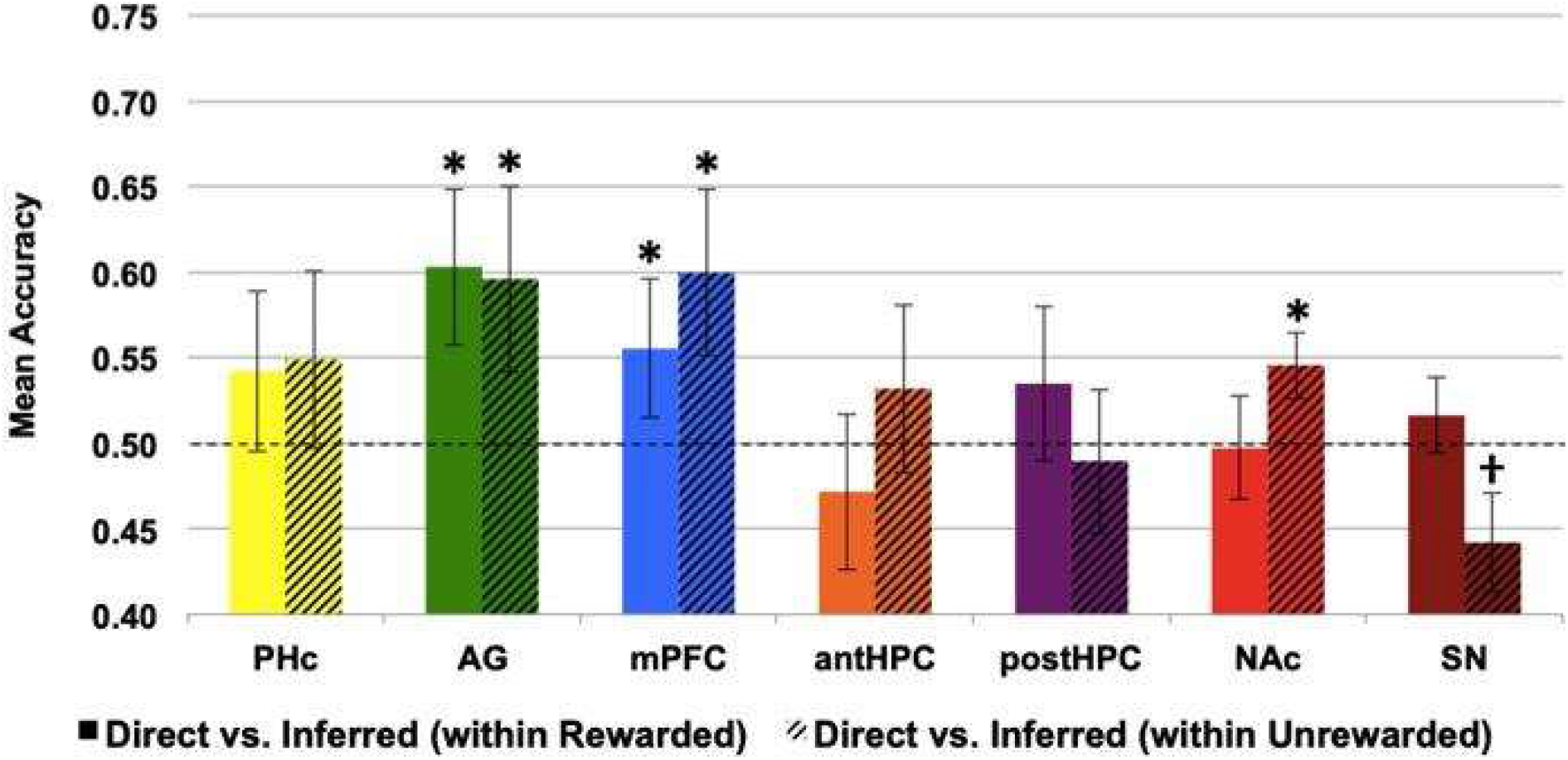
MVPA classifier accuracies for retrieval of rewarded (solid) and unrewarded (slashed) directly learned vs. inferred pairs (^*^ indicates significance, --- indicates chance). Error bars reflect standard error of the mean. Colors correspond to Figure 2

Finally, this process was repeated for retrieval of inferred and directly learned incongruent word pairs. Significantly distinct patterns were observed in the AG (M = .673, t(14) = 7.84, p < .001), mPFC (M = .623, t(14) = 3.66, p < .05), and posterior hippocampus (M = .572, t(14) = 2.31, p < .05), while the anterior hippocampus (M = .515, t(14) = .593, p = .567) and PHc (M = .520, t(14) = .926, p = .370) were not significantly above-chance (Fig. 11). An additional paired t-test of classifier accuracies for direct/inferred patterns within ROIs found significantly different accuracies for congruent and incongruent contrasts within the AG (t(14) = -2.60, p <.05) (Fig. 11).

Finally, MVPA was performed for rewarded vs. unrewarded items within directly learned and inferred pairs. Two-way repeated measures ANOVA was run between ROI and reward and found a main effect of ROI (F(6,84) = 6.662, p < 0.001, r =0.322), but no main effect of reward (F(1,14) = 0.136, p = 0.718, r = 0.010). Indicating that differences in classification for this contrast appeared due to differences between ROIs but not due to the influence of reward. While no effects were seen in the current study, additional investigation with a larger subject pool may have the power required to find effects of reward.

##### RSA

Representational Similarity Analysis (RSA) was conducted in order to determine if retrieval of linked AC pairs resulted in reactivation of its constituents (AB, BC). In a LME regression model for each ROI, connection type (AB-AC, BC-AC) was a significant predictor of the strength of pattern correlation in the PHc (B = -0.0382, t(57) = -2.728, p = 0.009), mPFC (B = -0.0374, t(57) = -2.035, p = 0.047), anterior hippocampus (B = -0.041, t(57) = -2.917, p = 0.005), and posterior hippocampus (B = -0.040, t(57) = -3.290, p = 0.002), but not in AG (B = - 0.028, t(57) = -1.278, p = .206). This suggests that the constituent reactivated during retrieval of linked pairs were important for all ROIs except the AG. Additionally, LME model results predicted that congruency was a significant predictor of the correlation between inferred pairs and their constituents in the PHc (B = -0.068, t(57) = -4.854, p < 0.001), AG (B = -0.044, t(57) = -2.009, p = 0.049), mPFC (B = -0.099, t(57) = -5.405, p < 0.001), anterior hippocampus (B = - 0.053, t(57) = -3.776, p < 0.001), and posterior hippocampus (B = -0.058, t(57) = -4.725, p < 0.001). Suggesting that correlations between linked pairs and their constituents were more similar for congruent sets than incongruent sets in all ROIs (see Fig. 13).

**Figure 13:**
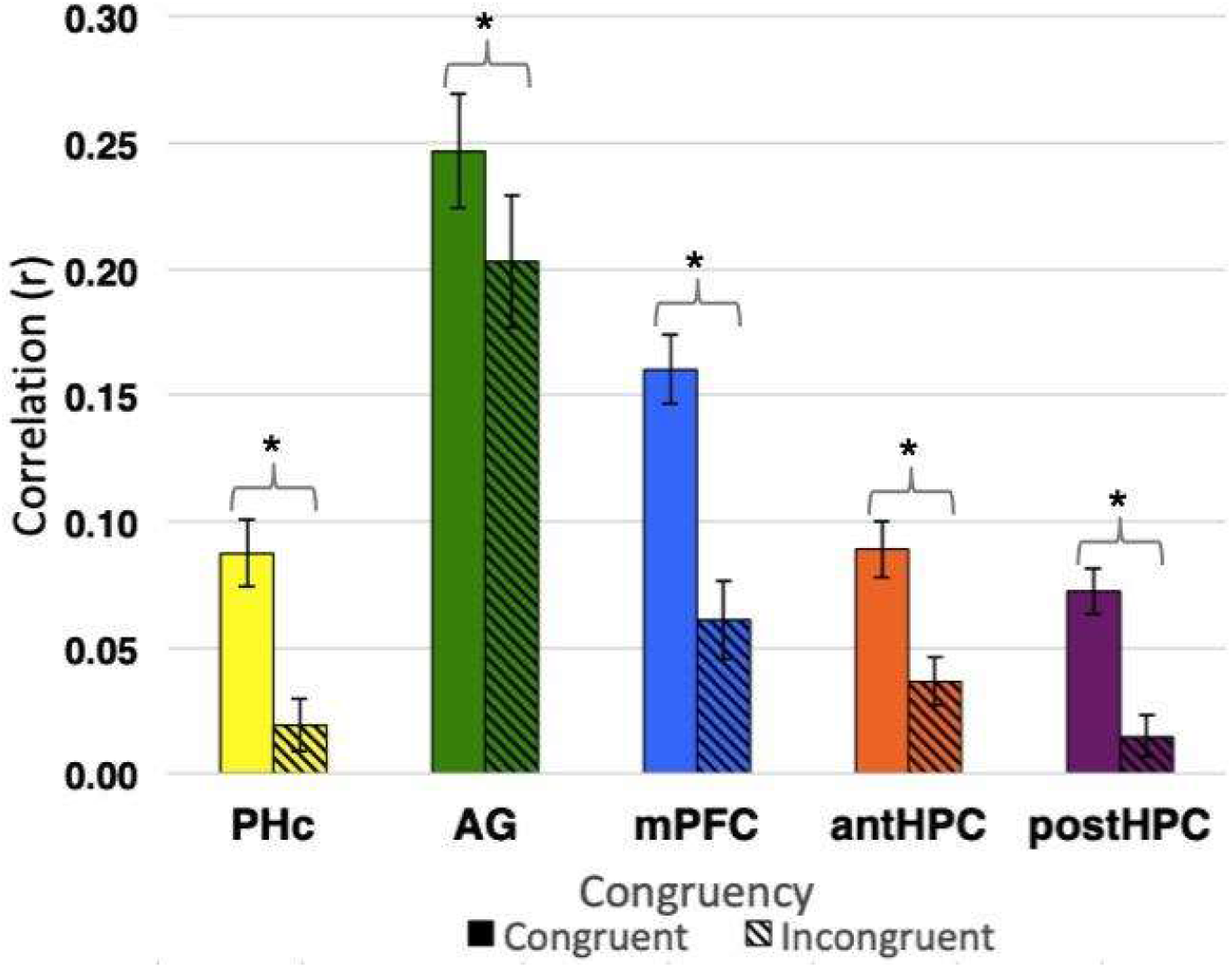
Correlations of inferred pair AC with its constituents AB and BC in schema congruent (solid) and incongruent (slashed) items (all bars showed significant differences between congruent and incongruent through a Linear Mixed Effects (LME) model). Error bars reflect standard error of the mean. Colors correspond to Figure 2

To investigate further similarities in neural patterns between inferred pairs and their constituents, inferred pairs in both congruent and incongruent sets were divided into two groups: successfully and unsuccessfully retrieved. Interestingly, a significant interaction between congruency and accuracy was observed in the mPFC (B = -0.076, t(955) = -2.053, p = 0.040), but was not seen in the PHc (B = 0.011, t(955) = 0.342, p = 0.733), AG (B = -0.067, t(955) = - 1.444, p = 0.149), anterior hippocampus (B = -0.008, t(955) = -0.241, p = 0.810), nor in the posterior hippocampus (B = 0.002, t(955) = 0.073, p = 0.942). These results show that differences in pattern similarity when pairs are successfully or unsuccessfully recalled is influenced by schema congruency in the mPFC (Fig. 14).

**Figure 14:**
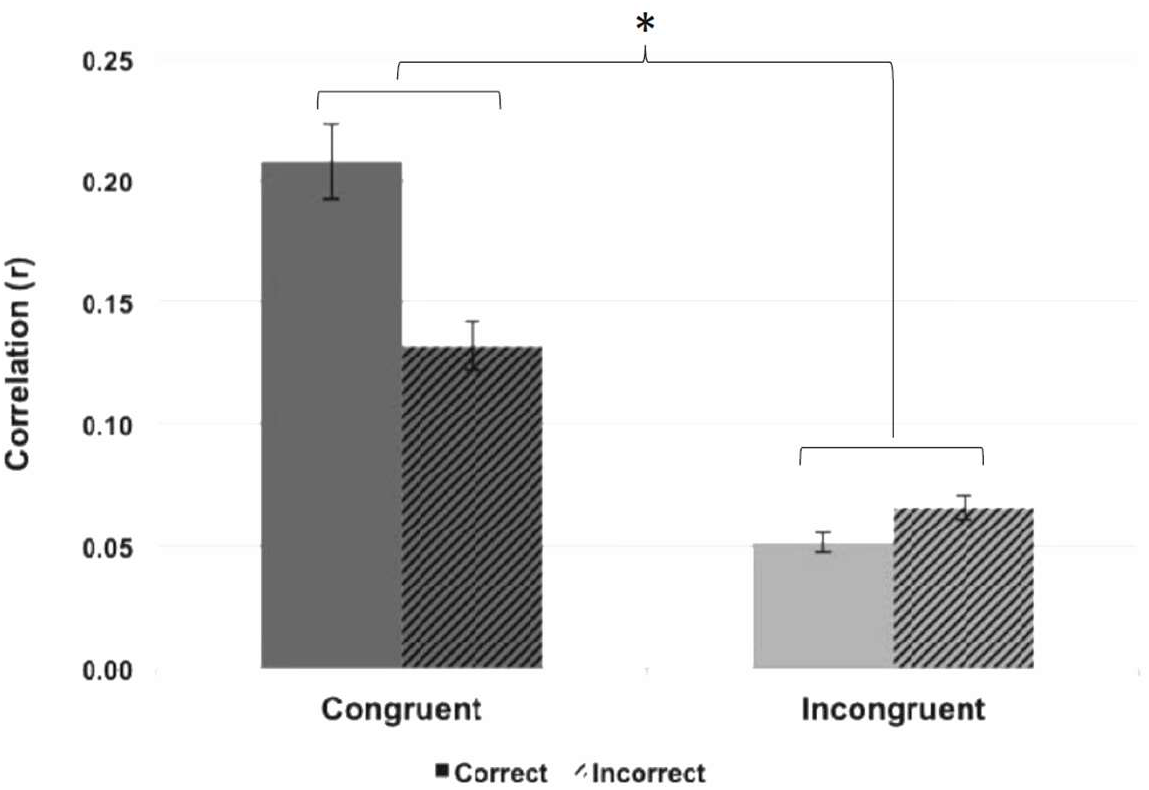
Correlations of inferred pair AC with its constituents AB and BC within the mPFC for correct (solid) and incorrect (slashed) congruent and incongruent pairs (bars show a significant interaction between accuracy and schema congruency through an LME model). Error bars reflect standard error of the mean.

### Between-Study Analysis

Finally, a two-way repeated measure mixed model ANOVA was conducted between MVPA results of directly learned vs. linked contrasts for experiment 1 and experiment 2, collapsed across reward, in order to determine if an interaction existed between rest and congruency. We found an interaction between rest and congruency (F(1,33) = 7.466, p = 0.010, r = 0.185).

Further, two-way repeated measures ANOVA between RSA correlation values for AB-AC/BC-AC between congruent and incongruent for experiment 1 and experiment 2, collapsed across reward, showed a significant interaction between sleep and congruency (F(1,35) = 8.451, p = 0.006, r = 0.194).

## Discussion

Using an associative inference task, we investigated how memory schema affects memory integration. After a night of sleep, the neural representations of schema-congruent inferred pairs were more similar to their constituent pairings (AB, BC), than schema-incongruent inferences, across regions of the memory network. Successfully integrated inferred pairs demonstrated higher pattern similarity in the mPFC with their directly learned constituents than did unsuccessfully integrated pairs, for congruent schema, but only after a night of sleep. These results suggest that delay between encoding and retrieval, paired with a delay with sleep, affects the influence of schema congruency on the brain’s memory network.

Behavioral measures confirmed that participants were able to successfully integrate both schema congruent and incongruent information regardless of the delay between encoding and retrieval. When retrieval immediately followed encoding, the accuracy of inferred pairs did not differ between congruent and incongruent sets; however, schema congruent inferred pairs were retrieved faster, suggesting that congruency may begin to have a beneficial effect on the integration of inferred pairs. Although not entirely conclusive due to lack of statistical significance in accuracy, these RT findings provide some support for prior results that associative pairs are better integrated when congruent with a schema (Zhang et al., 2018). When retrieval occurred the day following encoding, participants showed no difference in either response time or accuracy based on schema congruency. One possible explanation for this discrepancy could come from the experimental design of the training phase. In this study, the training task was modeled after the retrieval task, which itself was modified to be compatible for fMRI. Unlike the training phase used by Zhang et al. (2018) in which participants were initially shown the direct pairs with no foils and subsequently given a cued-recall test where they typed the target word, our training phase involved a forced-choice paradigm where the target word was always accompanied by a foil. It is possible that the presence of a foil during learning led to increased interference with the target word, which is consistent with the generally higher behavioral performance observed in Zhang et al. despite using similar stimuli and retrieval tasks.

Such interference would be especially detrimental for congruent pairs as they share semantic relationships, and could explain why we did not find such a strong effect of congruency.

Building on literature suggesting that associated items are represented more similarly than unrelated items during encoding (Schapiro et al., 2012), we investigated whether neural patterns for inferred pairs are represented similarly to the underlying learned pairs during retrieval. Additionally, we investigated whether factors that are known to improve memory performance such as sleep (Landmann et al., 2014, Stickgold and Walker, 2013) and reward-mediated motivation (Adcock et al., 2007, Knutson et al., 2001) would change the effect of congruency on memory integration. Using machine learning to decode multivariate patterns of activity, we identified regions that represented directly learned pairs differently from inferred pairs. In line with research suggesting that hippocampal and neocortical (Schlichting and Preston, 2014; Spalding et al., 2018) involvement is necessary for associative inference, we learned and inferred pairs were represented differently in the posHPC and mPFC. Additionally, the AG, which serves as a semantic hub for schema congruent information (Gilboa and Marlatte, 2017; Wagner et al., 2015), showed a similar effect. These findings were further supported by results in a second experiment, in which the mPFC and AG were again identified as representing direct and inferred pairs separately. The posHPC was no longer significant in this second experiment, possibly reflecting hippocampal disengagement as information is increasingly consolidated in the mPFC (van Kesteren et al., 2010a). Examining the role of schema congruency, surprisingly, the first experiment did not identify distinct neural representations for direct and inferred pairs, despite the benefits of schema congruency on memory (Zhang et al., 2018). Our second experiment revealed a difference in neural representations in the mPFC and AG between congruent and incongruent sets, with the latter showing increased pattern distinctiveness. A mixed-model repeated measures ANOVA between classifier accuracies from experiments 1 and 2 revealed a significant interaction between delay with sleep and congruency. These findings suggest that delay with sleep was necessary for the beneficial effect of schema congruency to be apparent.

The final examined modulation included in experiment 2 set out to investigate the influence of reward on memory integration and its interaction with schema. Behavioral data did not support our original hypothesis that reward would benefit both schema congruent and incongruent sets. Interestingly, retrieval performance of schema-congruent (but not schema-incongruent) pairs was hindered for rewarded word pairs that were congruent with a schema when they were associated with reward, while incongruent word pairs retrieval was not influenced by reward. One possible explanation is that reward led to increased interference between learned pairs and the foils of congruent sets. Evidence from prior literature suggests that reward reduces behavioral interference (Krebs et al., 2013), however disruptive effects of dopamine in the mPFC during working memory tasks have been observed in mice (Valentim Jr. et al., 2009), though reward has also reduced behavioral interference as well (Krebs et al., 2013). Increases in dopamine in the mPFC during reward-associated learning may have caused the behavioral detriment observed in this study. Additionally, increased activity of the mPFC during encoding and retrieval of schema congruent, but not incongruent, information (Brod et al., 2015, Ghosh et al., 2014) could explain why this was not observed in schema incongruent sets, which has been found to rely on greater hippocampal activity (van Kesteren et al., 2010a). Further investigation is required in order to fully understand the effect of reward on schematic memory. These early results in the examination of reward and schema suggest the two may not interact in this context, but further work is warranted to understand if this applies across all tasks and learned material.

RSA analysis that compared the neural representations of inferred pairs AC and their constituents (AB and BC) revealed greater pattern similarity for congruent than incongruent pairs in the PHc, mPFC, and posHPC. Subsequent analyses revealed that these results were not driven simply by shared semantic content between schema congruent pairs. These findings suggest that schema facilitates the assimilation of paired items into a single inferred unit containing all associated elements (Burton et al., 2016). In experiment 2, we found similar results, indicating that direct and inferred pairs were more similar for schema congruent pairs in the PHc, AG, mPFC, antHPC, and posHPC. Further analyses between both experiments revealed that a delay with sleep between encoding and retrieval increased the assimilation of inferred pairs. After a night of sleep, successfully integrated pairs were represented more similarly to their constituents than were unsuccessfully integrated pairs, in the mPFC, when the words were congruent with a schema. In incongruent sets, however, successful integration had no effect on pattern similarity. These findings expand on the prior literature of associative inference that has shown that reactivation of directly learned patterns during encoding of new events (i.e reactivation of AB when learning BC) predicts subsequent memory performance (Kuhl et al., 2010; Zeithamova et al., 2012). Our results suggest that the mPFC, which is heavily involved in schema (Brod et al., 2015), gradually shifts the representations of inferred pairs during retrieval to become more similar to direct pairs.

## Acknowledgements

The authors thank the University of Pittsburgh and Carnegie Mellon University Brain Imaging Data Generation & Education (BRIDGE) Center for funding the second experiment of this study. We also thank Griffin Koch, Heather Bruett, and Jasmine Issa for contributing to the creation of stimuli, and Mark Vignone and Scott Kurdilla for their assistance in running subjects. *Conflict of Interest*: The authors declare no competing financial interests.

